# Extensive Ribosome and RF2 Rearrangements during Translation Termination

**DOI:** 10.1101/600445

**Authors:** Egor Svidritskiy, Gabriel Demo, Anna B. Loveland, Chen Xu, Andrei A. Korostelev

## Abstract

Protein synthesis ends when a ribosome reaches an mRNA stop codon. Release factors (RFs) decode the stop codon, hydrolyze peptidyl-tRNA to release the nascent protein, and then dissociate to allow ribosome recycling. To visualize termination by RF2, we resolved a cryo-EM ensemble of *E. coli* 70S•RF2 structures at up to 3.3 Å in a single sample. Five structures suggest a highly dynamic termination pathway. Upon peptidyl-tRNA hydrolysis, the CCA end of deacyl-tRNA departs from the peptidyl transferase center. The catalytic GGQ loop of RF2 is rearranged into a long β-hairpin that plugs the peptide tunnel, biasing a nascent protein toward the ribosome exit. Ribosomal intersubunit rotation destabilizes the catalytic RF2 domain on the 50S subunit and disassembles the central intersubunit bridge B2a, resulting in RF2 departure. Our structures visualize how local rearrangements and spontaneous inter-subunit rotation poise the newly-made protein and RF2 to dissociate in preparation for ribosome recycling.

## Introduction

Ribosomes terminate translation when release factors (RFs) recognize an mRNA stop codon in the A site (Brenner et al., 1967; Brenner et al., 1965). Upon stop-codon recognition, the release factor hydrolyzes peptidyl-tRNA in the P site of the large subunit, thus releasing nascent protein from the ribosome (Capecchi, 1967a, b; Scolnick et al., 1968; Tompkins et al., 1970). In bacteria, RF1 recognizes UAA or UAG, and RF2 recognizes UAA or UGA codons (Nakamura et al., 1995). Following peptide release, RF1 or RF2 must dissociate from the ribosome to allow the ribosome and release factor to recycle. While structural studies have provided snapshots of some peptidyl-tRNA hydrolysis steps (reviewed in (Dunkle and Cate, 2010; Korostelev, 2011; Ramakrishnan, 2011; Rodnina, 2018; Youngman et al., 2008), recent biophysical and biochemical findings suggest a highly dynamic series of termination events (Adio et al., 2018), which have not been structurally visualized.

In the pre-termination (pre-hydrolysis) state, the substrate peptidyl-tRNA resides in a nonrotated (classical) conformation of 70S ribosome, with its anticodon stem loop bound to the mRNA codon in the P site of the small 30S subunit and the CCA-peptidyl moiety in the P site of the large 50S subunit (Agrawal et al., 2000; Ermolenko et al., 2007). Cryogenic electron microscopy (cryo-EM) and X-ray studies reveal RF1 or RF2 bound in the A site next to P tRNA (Klaholz et al., 2003; Petry et al., 2005; Rawat et al., 2006; Rawat et al., 2003), and provide insight into pre-hydrolysis (Jin et al., 2010) and post-hydrolysis states (Korostelev et al., 2008; Korostelev et al., 2010; Laurberg et al., 2008; Weixlbaumer et al., 2008). The release factor’s codon-recognition domain binds the stop codon in the 30S decoding center. The catalytic GGQ motif inserts into the 50S peptidyl-transferase center (PTC) adjacent to the CCA end of P-tRNA. Ribosomes retain the non-rotated conformation in RF-bound structures, including those with mutated or post-translationally modified RFs (Pierson et al., 2016; Santos et al., 2013; Zeng and Jin, 2018) or when RFs are perturbed by the termination inhibitor blasticidin S (Svidritskiy and Korostelev, 2018a, b) or by a sense codon in the A site (Svidritskiy et al., 2018). The catalytic GGQ loop adopts the same compact conformation comprising a short α-helix in pre-hydrolysis-like and post-hydrolysis structures, as discussed below.

Ribosome and RF dynamics are implicated in post-hydrolysis steps. In *E. coli,* RF1 dissociation is assisted by the non-essential GTPase RF3 (Freistroffer et al., 1997; Koutmou et al., 2014), whereas RF2 can spontaneously dissociate from the ribosome (Adio et al., 2018). RF3 induces rotation of the small subunit relative to the large subunit by ~9 degrees in the absence of RF1 and RF2 (Ermolenko et al., 2007; Zhou et al., 2012) and stabilizes a “hybrid” state of deacylated tRNA (P/E) (Gao et al., 2007; Jin et al., 2011). In the P/E conformation, the anticodon stem loop remains in the 30S P site, while the elbow and acceptor arm bind the L1 stalk and E site of the 50S subunit. A recent cryo-EM study (Graf et al., 2018) showed that if RF1 is locked on the post-hydrolysis ribosome by antibacterial peptide apidaecin 137 (Florin et al., 2017), the ribosome can undergo intersubunit rotation in the presence of both RF1 and RF3. By contrast, biochemical experiments showed that locking RF1 on the ribosome by inactivating its catalytic motif (GGQ-> GAQ) prevented RF1 recycling by RF3 from the pre-hydrolysis ribosome (Koutmou et al., 2014). These and other (Adio et al., 2018) observations indicate that peptidyl-tRNA hydrolysis is critical for RF1 dissociation, likely due to propensity of deacylated tRNA for exiting the P site and sampling the P/E hybrid state via spontaneous intersubunit rotation (Moazed and Noller, 1989). Nevertheless, RF3 is dispensable for growth of *Escherichia coli* (Grentzmann et al., 1994; Mikuni et al., 1994), and its expression is not conserved in bacteria (Leipe et al., 2002). For example, RF3 is not present in the thermophilic model organisms of the *Thermus* and *Thermatoga* genera and in infectious *Chlamydiales* and *Spirochaetae.* This means that both RF1 and RF2 are capable of performing a complete round of termination independently of RF3. Indeed, recent Förster Resonance Energy Transfer (FRET) studies demonstrate spontaneous RF1 and RF2 dissociation after peptidyl-tRNA hydrolysis (Prabhakar et al., 2017), coupled with transient RF sampling of the rotated ribosomes (Adio et al., 2018). These observations, however, appear to contradict cryo-EM, X-ray crystallographic and FRET studies (Casy et al., 2018; Sternberg et al., 2009) that suggest RF1 and RF2 stabilize the nonrotated 70S conformation. Thus it remains unclear how RFs alone terminate translation and dissociate from the ribosome, and whether these steps involve ribosome rearrangements.

In this work, we used ensemble cryo-EM (Abeyrathne et al., 2016; Loveland et al., 2017) to visualize the structural dynamics of RF2 on the 70S ribosome (**Figure 1, Figure 1—figure supplement 1 and Figure 1—figure supplement 2**). The structures capture RF2 and tRNA in distinct 70S conformational states (Table 1). They reveal an unexpected rearrangement of the catalytic GGQ loop, which suggests that RF2 helps direct nascent protein out of the ribosome. Our findings reconcile biophysical observations and, together with other structural studies, allow a reconstruction of the termination course by RF2, from initial binding to dissociation.

**Figure 1.**
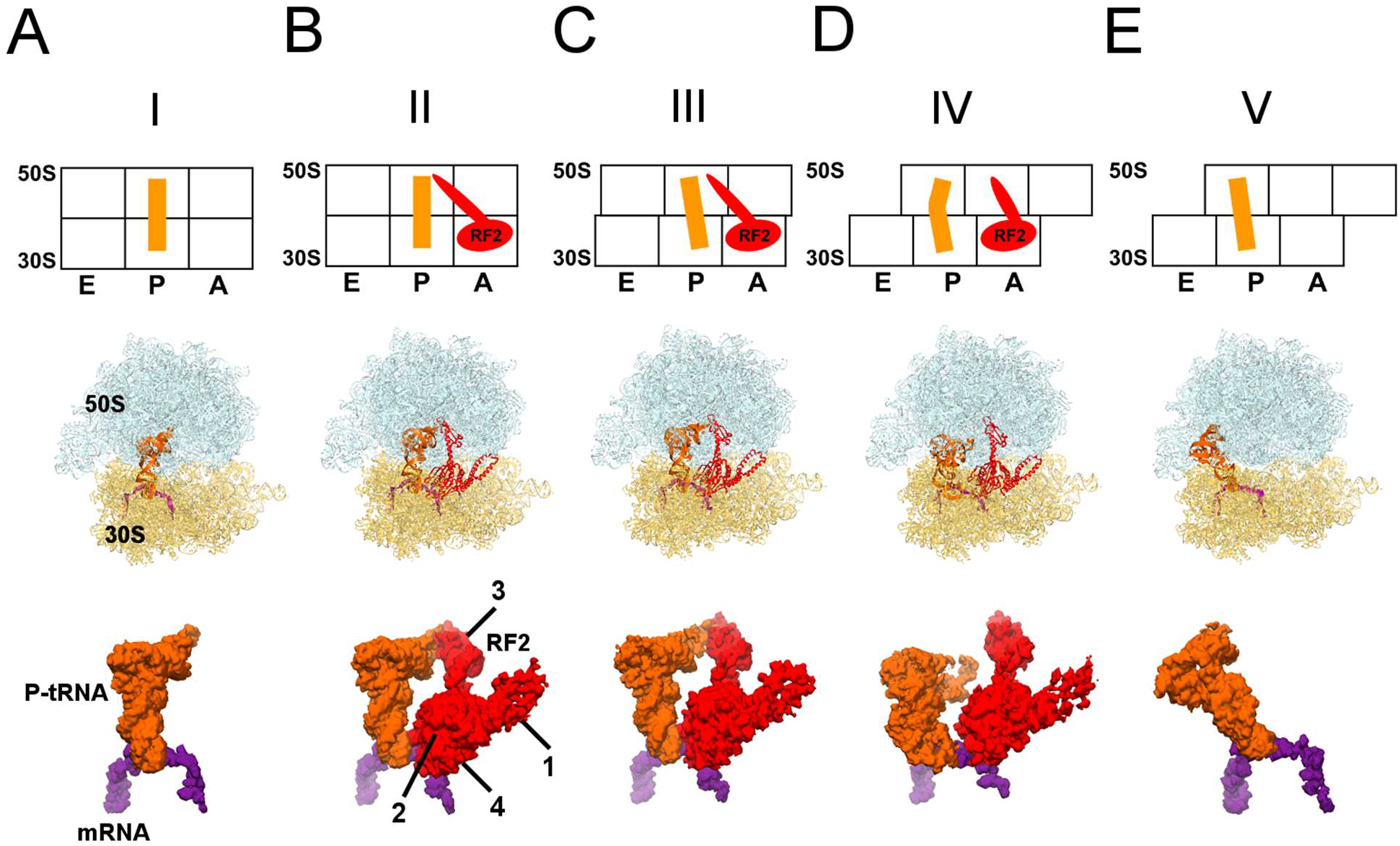
Cryo-EM Structures I to V. Panels A-E show for each structure: *upper row: a* schematic of the conformations of the 70S ribosome, tRNA and RF2; *middle row:* structures with RF2 shown in red, tRNA^fMet^ in orange, 30S subunit in yellow, 50S subunit in cyan and mRNA in purple; *lower row:* cryo-EM density for mRNA, tRNA and RF2, colored as in the middle row.

**Table 1.** Cryo-EM structures obtained from a single sample of the 70S•RF2 complex.

|  | I | II | III | IV | V |
| --- | --- | --- | --- | --- | --- |
| **tRNA and RF2 occupancy** | P-tRNA | P-tRNA, RF2 | P-like (L5) tRNA, RF2 | P/E-like (L1) tRNA, RF2 | P/E tRNA |
| **30S (body) rotation** | 1.6° | 1.6° | 4.9° | 7.4° | 6.8° |
| **30S head rotation (swivel)** | 2.0° | 1.4° | 1.7° | 2.8° | 6.0° |
| **Resolution (FSC=0.143)** | 3.3 Å | 3.3 Å | 3.9 Å | 4.2 Å | 3.4 Å |

## Results and Discussion

We formed a termination complex using *E. coli* 70S ribosomes and RF2 to catalyze the hydrolysis of substrate N-formyl-methionyl-tRNA^fMet^ (fMet-tRNA^fMet^) in the presence of mRNA carrying an AUG codon in the P site and a UGA stop codon in the A site. Termination complexes for cryo-EM were formed under conditions that support hydrolysis of fMet-tRNA^fMet^ and release of fMet (**Figure 1—figure supplement 3**), demonstrating activity of RF2 in our sample. Maximum-likelihood classification of cryo-EM data in FREALIGN (Lyumkis et al., 2013) yielded five structures at average resolutions of 3.3 Å to 4.2 Å (Table 2; **Figure 1**). The structures differ by the degrees of intersubunit rotation and 30S head rotation (swivel), and are consistent with the following termination stages: pre-termination (i.e., no RF2; Structure I); peptidyl-tRNA hydrolysis by RF2 (Structure II); tRNA exit from the PTC (Structure III); destabilization of the catalytic domain of RF2 (Structure IV); and post-termination (i.e., after RF2 dissociates; Structure V).

**Table 2.**
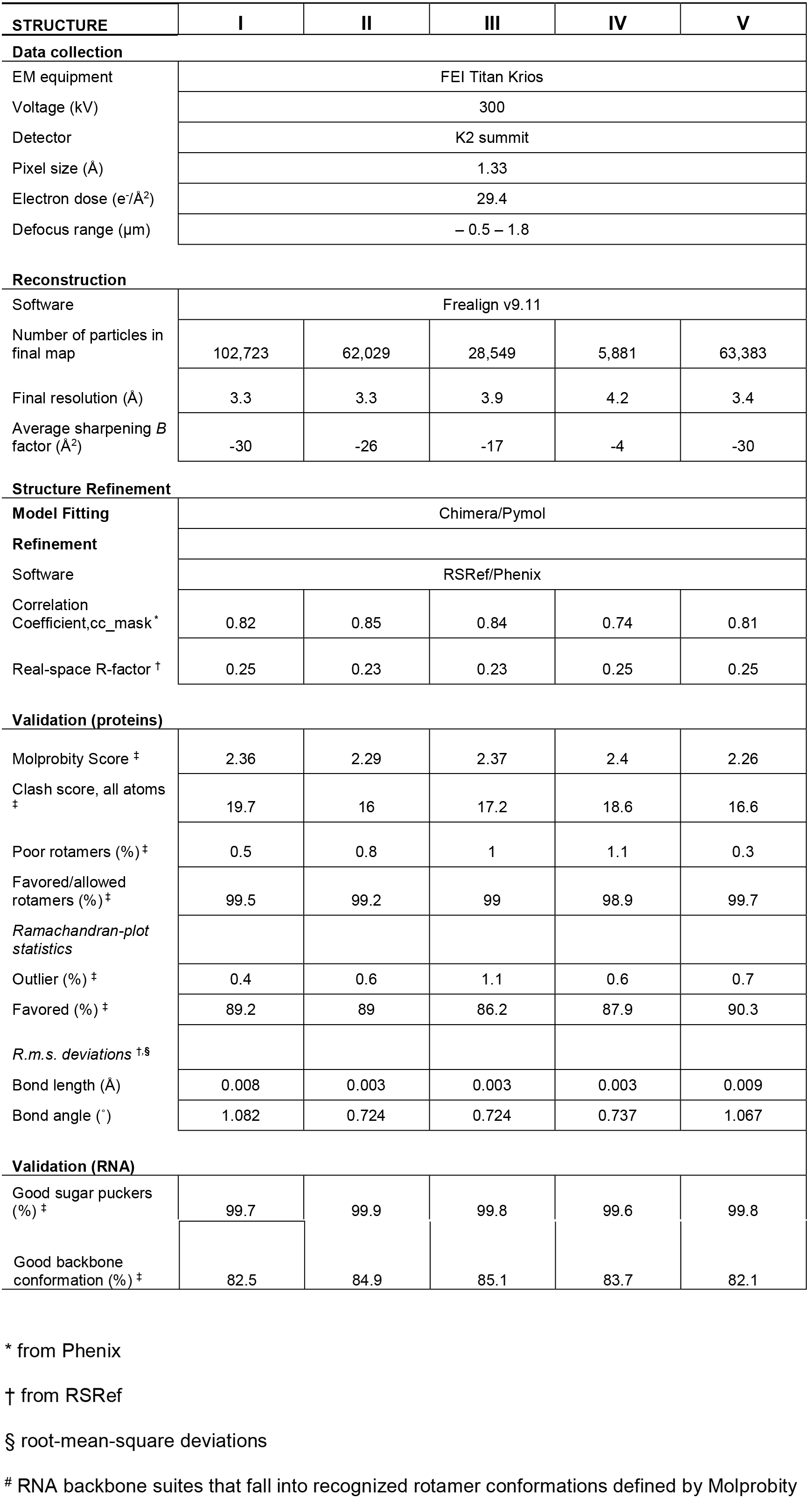
Data collection and refinement statistics for cryo-EM Structures I-V.

Pre-termination Structure I contains tRNA in the P site with its 3’-CCA end positioned in the peptidyl transferase center, similar to that in aminoacyl- and peptidyl-tRNA complexes (**Figure 1A**). The 30S subunit adopts a conformation similar to crystal structures of non-rotated ribosomes with or without RFs (Jin et al., 2010; Korostelev et al., 2008; Korostelev et al., 2006; Korostelev et al., 2010; Laurberg et al., 2008; Selmer et al., 2006; Weixlbaumer et al., 2008), and exhibits a slight ~1.5° rotation relative to the 70S•RF2 crystal structure formed with the UGA stop codon (Weixlbaumer et al., 2008).

RF2 is bound in Structures II to IV (**Figures 1B-D**). 70S intersubunit rotation ranges from ~1.5° in Structure II to ~7.5° in Structure IV (Table 1). This nearly full range of rotation is typical of translocation complexes with tRNA (Agirrezabala et al., 2008; Agirrezabala et al., 2012; Dunkle et al., 2011; Fischer et al., 2010; Julian et al., 2008) and elongation factor G (EF-G) (Brilot et al., 2013; Chen et al., 2013; Frank and Agrawal, 2000; Gao et al., 2009; Pulk and Cate, 2013; Tourigny et al., 2013; Zhou et al., 2013).

The most well-resolved 3.3-Å resolution Structure II (**Figure 1B**) accounts for the largest population of RF2-bound ribosome particles in the sample (Table 2), consistent with stabilization of the non-rotated 30S states by RF2 observed in FRET studies (Casy et al., 2018; Prabhakar et al., 2017; Sternberg et al., 2009). The overall extended conformation of RF2 resembles that seen in published 70S•RF2 structures (Korostelev et al., 2008; Weixlbaumer et al., 2008) with its codon-recognition superdomain (domains 2 and 4) bound to the UGA stop codon at the decoding center of the 30S subunit and its catalytic domain 3 inserted into the peptidyl-transferase center.

Structure II features RF2 with a dramatically rearranged catalytic loop, coinciding with the lack of density for the CCA end of the P tRNA. In previous pre- and post-hydrolysis-like structures, the P-tRNA CCA end binds the PTC, and RF2 forms a compact catalytic loop (residues 245-258). The catalytic ^250^GGQ^252^ motif resides at the tip of a short α-helix, adjacent to the terminal tRNA nucleotide A76 (**Figure 2A**, light shades; (Jin et al., 2010; Korostelev et al., 2008; Laurberg et al., 2008; Weixlbaumer et al., 2008)). The α-helical region is held in place by A2602 of 23S rRNA, which is critical for termination efficiency (Amort et al., 2007; Polacek et al., 2003). In Structure II, however, the catalytic loop adopts an extended β-hairpin conformation that reaches ~10 Å deeper into the peptide tunnel (**Figures 2A-B**). To accommodate the extended β-hairpin, nucleotides G2505 and U2506 of 23S rRNA are shifted to widen the PTC (**Figure 2— figure supplement 1A**). The center of the β-hairpin occupies the position normally held by A76 of P tRNA (**Figure 2A**). The lack of density for the CCA moiety thus suggests that the 3’ end of P tRNA is released from the PTC. Indeed, formation of the β-hairpin and release of the CCA moiety is accompanied by a slight shift in the P-tRNA acceptor arm away from the PTC and by ~160° rotation of nucleotide A2602 to form a Hoogsteen base pair with C1965 at helix 71 of 23S rRNA (**Figure 2A and Figure 2—figure supplement 1B**). The cryo-EM map shows strong density for the extended β strands (**Figure 2C**), but weaker density for the GGQ residues at the tip of the hairpin, consistent with conformational heterogeneity of the flexible glycine backbone. Nevertheless, the tip of the β-hairpin plugs the narrowest region of the peptide tunnel at A2062 of 23S rRNA (**Figure 2B**). Thus, Structure II appears to represent a previously unseen post-peptide-release state with deacyl-tRNA dissociating from the PTC.

**Figure 2.**
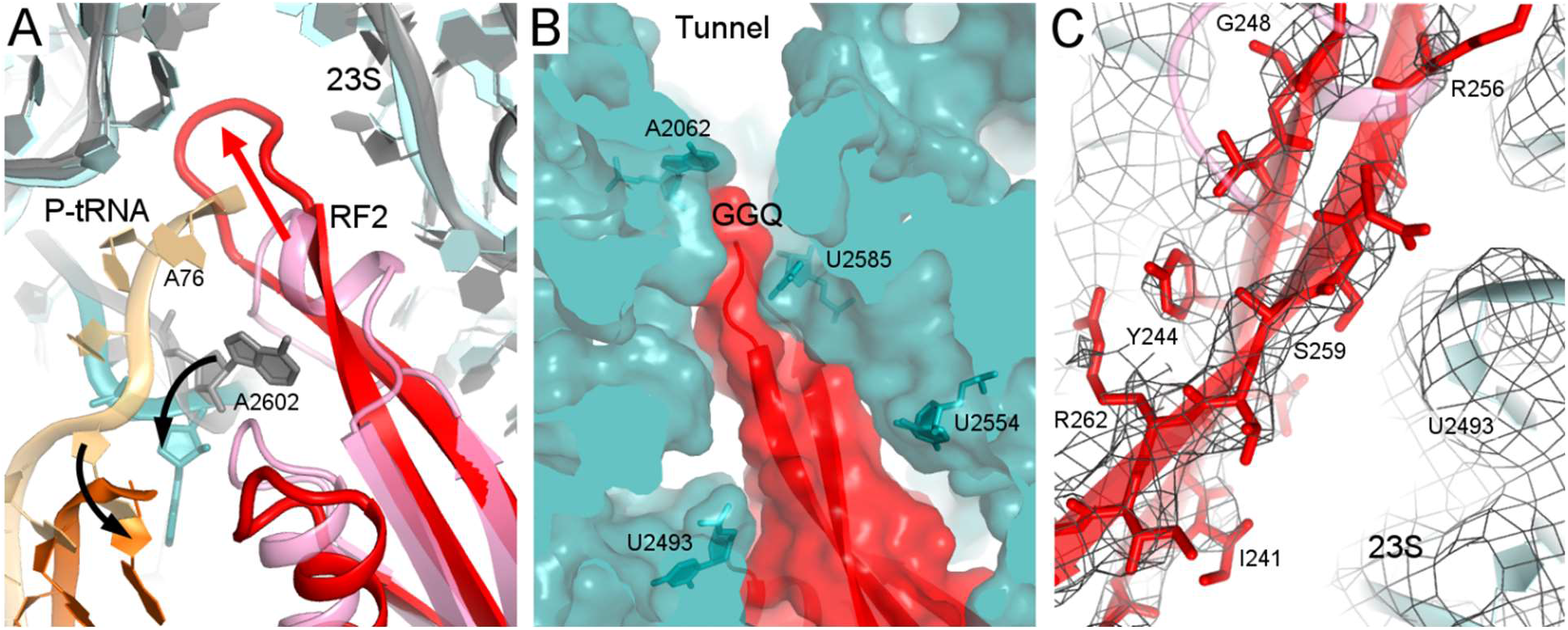
Rearrangement of the catalytic domain of RF2 upon tRNA displacement from the PTC. **A**. Rearrangements of RF2 in Structure II (RF2 in red, P-tRNA in orange and 23S in cyan) in comparison with the canonical 70S•RF2 conformation (X-ray structure: (Korostelev et al., 2008); RF2 in magenta, P-tRNA in light orange, 23S in gray). Arrows show rearrangements for RF2 (red), acceptor arm (orange) and 23S nucleotide A2602 (teal). The superposition was achieved by structural alignment of 23S rRNA. **B**. The GGQ region of RF2 forms a long β-hairpin that reaches into the constriction of the peptide tunnel (at A2062) and thus plugs the tunnel. Nucleotides of 23S rRNA and the GGQ motif (gray) are labeled. **C**. Cryo-EM density for the β-hairpin formed by the GGQ region. Residues of RF2 and 23S rRNA (also shown in panel B for reference) are labeled.

In Structure III (**Figure 1C**), a 5° rotation of the 30S subunit shifts the tRNA elbow by ~10 Å along protein L5 toward the E site. As a result, the P-tRNA acceptor arm pulls further out of the 50S P site, and the end of the acceptor arm helix points at helix α7 of RF2 near positively charged Lys 282 and Lys 289. The CCA moiety remains unresolved. The 5° intersubunit rotation also causes a slight 2° rotation of RF2 domain 3 relative to domain 2, but the overall conformation of RF2 remains similar to that in Structure II, with the tip of the extended β-hairpin plugging the peptide exit tunnel.

In Structure IV, the 30S subunit is rotated 7.5° and the P-tRNA elbow is bound to the L1 stalk, an interaction normally observed with the P/E or E tRNA (Dunkle et al., 2011; Korostelev et al., 2006; Selmer et al., 2006). Unlike P/E tRNA, however, the acceptor arm is oriented between the 50S P and E sites (**Figure 1D**), suggesting that the tRNA adopts a transient state on its way toward the hybrid P/E conformation. This transition coincides with an ~3° rotation (swivel) of the 30S head, characteristic of intermediate stages of tRNA and mRNA translocation (Abeyrathne et al., 2016; Ratje et al., 2010). The codon-recognition superdomain of RF2 in its canonical conformation moves with the 30S subunit, causing the catalytic domain to shift by ~5 Å from its position in Structure II (**Figure 3A**). Indeed, reduced density for the catalytic domain and domain 1, which bridges the 30S and 50S subunits at the A site periphery (**Figure 1**), suggests that intersubunit rotation has destabilized RF2. In Structure IV, therefore, RF2 appears to be poised for dissociation and collapsing toward the compact conformation adopted by free RFs (**Figure 3A**) (JCSG, 2005; Shin et al., 2004; Vestergaard et al., 2001; Zoldak et al., 2007).

**Figure 3.**
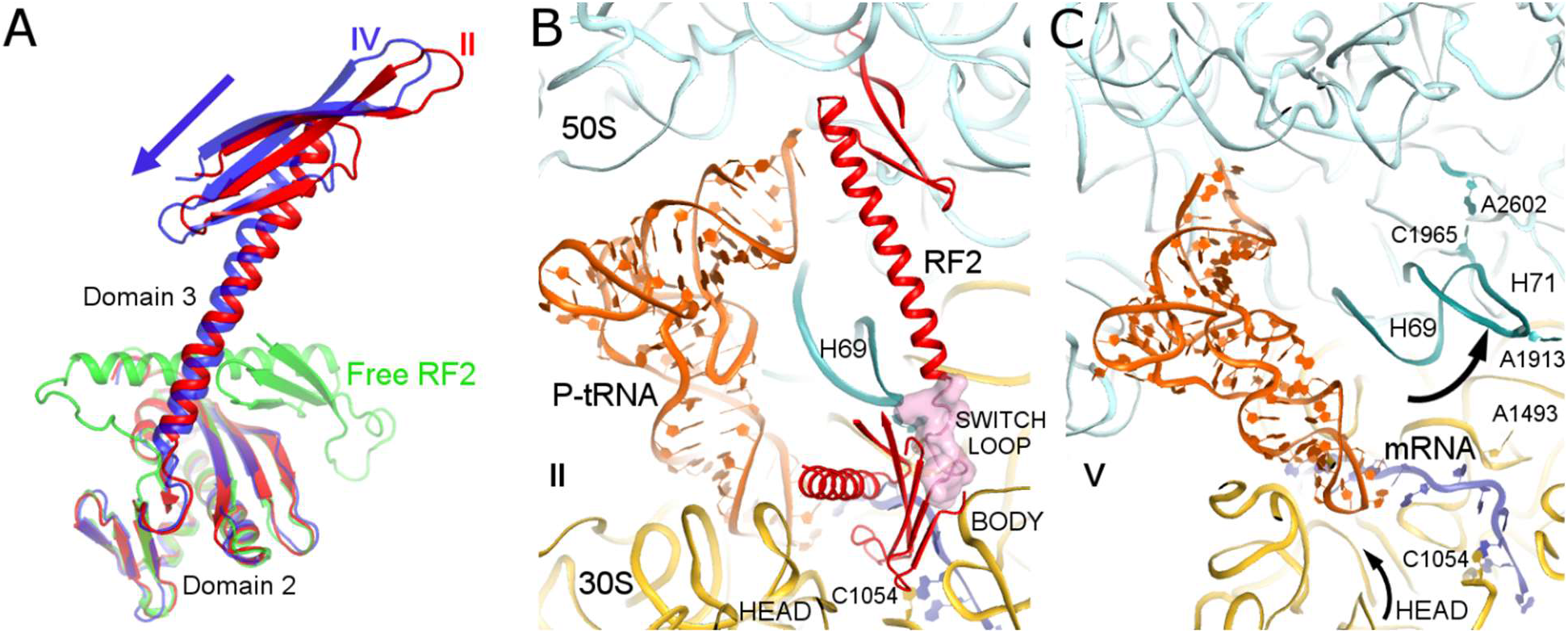
Differences in conformations of RF2 and helix 69 of 23S rRNA in Structures II, IV and V. **A**. RF2 in Structures II (red) and IV (blue) in comparison with the crystal structure of isolated (free) RF2 (green, PDB 1GQE (Vestergaard et al., 2001)). The arrow shows the direction of domain 3 repositioning from Structure II to IV to free RF2. Structures were aligned by superposition of domain 2. **B**. Interactions of H69 (teal) with the switch loop (pink surface) of RF2 in Structure II (RF2 in red, P-tRNA in orange, 50S in cyan and 30S in yellow). Domains 2 and 3 of RF2, head and body domains of the 30S subunit and nucleotides A1913 (at H69) and C1054 (head) are labeled. **C**. In Structure V, dissociation of H69 from the decoding center and packing on H71 next to A2602 disassembles intersubunit bridge B2a. The structure is colored as in panel B. View in panels B and C is rotated by ~180° relative to panel A.

Structure V is incompatible with RF2 binding and provides further insights into RF2 dissociation. The rotated 70S•tRNA structure resembles hybrid P/E states, with the acceptor arm of tRNA directed toward the E site of 50S (**Figure 1E**). However, the central intersubunit bridge B2a— formed by helix 69 (H69, nt. 1906-1924) of 23S rRNA—is disassembled (**Figures 3B-C**). In RF-bound structures (including Structures I to IV, above), the tip of helix 69 (at nt 1913) locks into the decoding center in a termination-specific arrangement (Korostelev et al., 2008; Korostelev et al., 2010; Laurberg et al., 2008; Weixlbaumer et al., 2008). H69 interacts with the switch loop of RFs, where the codon-recognition superdomain connects to catalytic domain 3 (e.g., Structure II, **Figure 3B**). This interaction directs the catalytic domain toward the PTC (Korostelev et al., 2010; Laurberg et al., 2008) and thus defines the efficiency and accuracy of release factors (Svidritskiy and Korostelev, 2018a). In Structure V, however, H69 is disengaged from the decoding center and moved ~15 Å toward the large subunit to pack against H71 near A2602 (nt 1945-1961; **Figure 3C**). The new position of H69 would clash with extended or compact conformations of RF2 (**Figure 3—figure supplement 1**), if the release factor had remained bound to the decoding center.

Release of RF2 also appears to be coupled to rotation (swivel) of the 30S head domain. In RF2-bound structures, the conserved codon-recognition SPF motif of RF2 interacts with the stop codon and the 30S head near bulged nucleotide C1054 of 16S rRNA. In Structure V, a 6° rotation of the 30S head shifts the position of C1054 by 5 Å from its position in RF2-bound Structure IV, consistent with disassembly of RF2 contacts. The disruption of decoding-center interactions due to repositioning of H69 and head swivel therefore explain the absence of RF2 in Structure V.

### Mechanism of Translation Termination by RF2

Our termination structures allow us to reconstruct a dynamic pathway for termination (**Figure 4 and Movie 1**) and reconcile biophysical and biochemical findings (Adio et al., 2018; Casy et al., 2018; Prabhakar et al., 2017; Sternberg et al., 2009). Recent structural studies revealed compact RF conformations on non-rotated 70S ribosomes (Fu et al., 2018; Svidritskiy and Korostelev, 2018a) that suggest how RFs open at early stages of codon recognition (He and Green, 2010; Hetrick et al., 2009; Trappl and Joseph, 2016). The short-lived transition from compact to open conformation(s) lasts tens of microseconds (Fu et al., 2018), and compact RF2 conformations were not captured in our sample. Compact RF2 was also visualized by cryo-EM on the 70S ribosome in the presence of truncated mRNA and alternative rescue factor A (ArfA) (Demo et al., 2017; James et al., 2016), suggesting a conserved pathway for initial binding of RFs to non-rotated ribosomes. Together with these recent studies, our work provides an expanded structural view of the termination course – from initial codon recognition to peptide release and RF2 dissociation (**Figure 4**).

**Figure 4.**
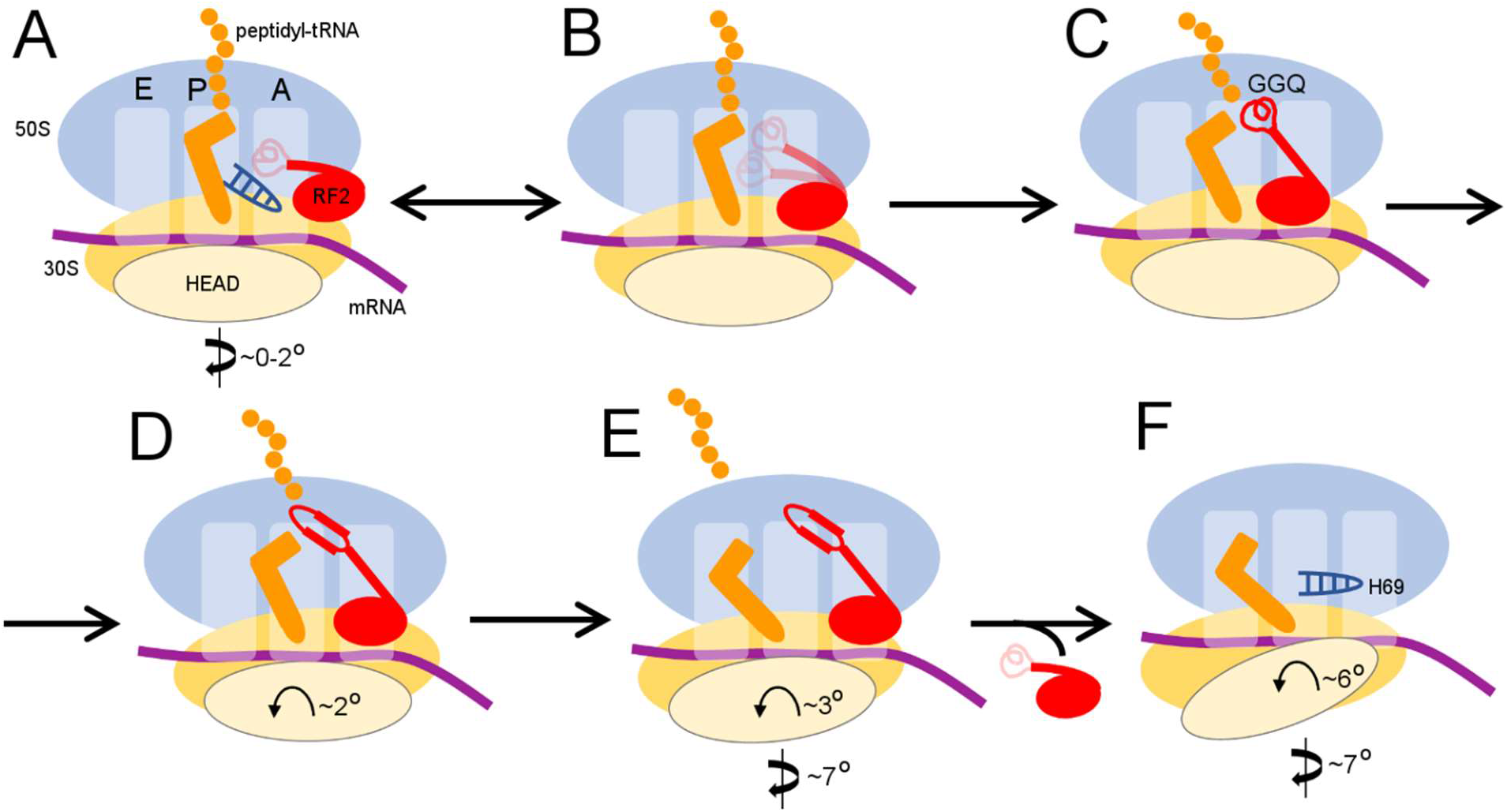
Scheme of the termination mechanism.

Termination begins when release factor recognizes a stop codon in the A site of a non-rotated (or slightly rotated) 70S ribosome carrying peptidyl-tRNA (**Figure 4A**). A compact release factor recognizes the stop codon, while its catalytic domain is kept 60 to 70 Å away from the PTC (Fu et al., 2018). Separation of the codon-recognition and catalytic functions during initial binding of RF underlies high accuracy of termination (Svidritskiy and Korostelev, 2018a). Upon binding the decoding center, steric hindrance with P-tRNA forces the RF catalytic domain to undock from domain 2 and sample the intersubnit space (**Figure 4B**) (discussed in (Svidritskiy and Korostelev, 2018a)). The ribosome remains non-rotated because peptidyl-tRNA bridges the P sites of 30S and 50S subunits. Interactions between the switch loop of RF and the decoding center and H69 of 30S direct the catalytic domain into the PTC. The GGQ catalytic loop is placed next to the scissile ester bond of peptidyl-tRNA and stabilized by A2602 (Jin et al., 2010; Laurberg et al., 2008), so that the Gln backbone amide of the GGQ motif can catalyze peptidyl-tRNA hydrolysis (**Figure 4C**) (Korostelev et al., 2008; Santos et al., 2013). These ribosome and RF2 rearrangements complete the first stage of termination leading to peptidyl-tRNA hydrolysis—an irreversible step that separates the newly made protein from P-site tRNA and prevents further elongation.

To be recycled, the ribosome must release the nascent protein and RF. The deacylated CCA end of the tRNA exits the 50S P site, which allows movement of A2602. No longer stabilized by the tRNA and A2602, the GGQ loop reorganizes into an extended β-hairpin that blocks the peptide tunnel (Structure II), thus biasing the diffusion of nascent peptide toward the exit of the peptide tunnel at the surface of the 50S subunit (**Figure 4D**). Furthermore, the exit of the CCA end from the P site supports formation of the P/E-tRNA hybrid conformation via spontaneous intersubunit rotation (Structures III and IV), which is also essential for translocation of tRNA-mRNA (reviewed in (Ling and Ermolenko, 2016; Noller et al., 2017)). The large counterclockwise rotation of the small subunit destabilizes the catalytic domain of RF2 (Structure IV), so that it begins to collapse toward domain 2 bound to the 30S subunit (**Figure 4E**). Rotation of the 30S head and detachment of H69 from the decoding center force dissociation of RF2 from the A site of the 30S subunit (Structure V, **Figure 4F**).

Remarkably similar extents of intersubunit rotation and head rotation were observed in recent RF1-bound 70S structures formed with RF3•GDPNP in the presence of apidaecin (which stalls RF1 on the ribosome). In the presence of stalled RF1, however, the intersubunit bridge remains attached to the decoding center (Graf et al., 2018). Thus the ribosome samples similar rotation states with RF1 and RF2, with or without RF3. RF3 stabilizes the rotated state (Ermolenko et al., 2007; Gao et al., 2007; Jin et al., 2011; Zhou et al., 2012) to promote RF release. However, the role of RF3 in termination is dispensable because intersubunit rotation is an inherent— i.e., spontaneous and thermally driven—property of the ribosome (Cornish et al., 2008). Cold sensitivity of RF3-deletion strains (Nichols et al., 2011; O’Connor, 2015) perhaps manifests the dependence on RF3 due to perturbation of the intersubunit dynamics at lower temperatures. In the fully rotated state, the post-termination ribosome becomes a substrate for subunit dissociation by EF-G and recycling factor RRF (Agrawal et al., 2004; Dunkle et al., 2011; Fu et al., 2016; Prabhakar et al., 2017). Structure V is consistent with structural studies showing ribosome recycling factor (RRF) bound to ribosomes with a dislodged intersubunit bridge B2a (Borovinskaya et al., 2007; Pai et al., 2008), which helps split the ribosome into subunits. Translation termination and recycling of the release factors and the ribosome therefore rely on the spontaneous ribosome dynamics, triggered by local rearrangements of the universally conserved elements of the peptidyl-transferase and decoding centers.

**Figure 1—figure supplement 1.**
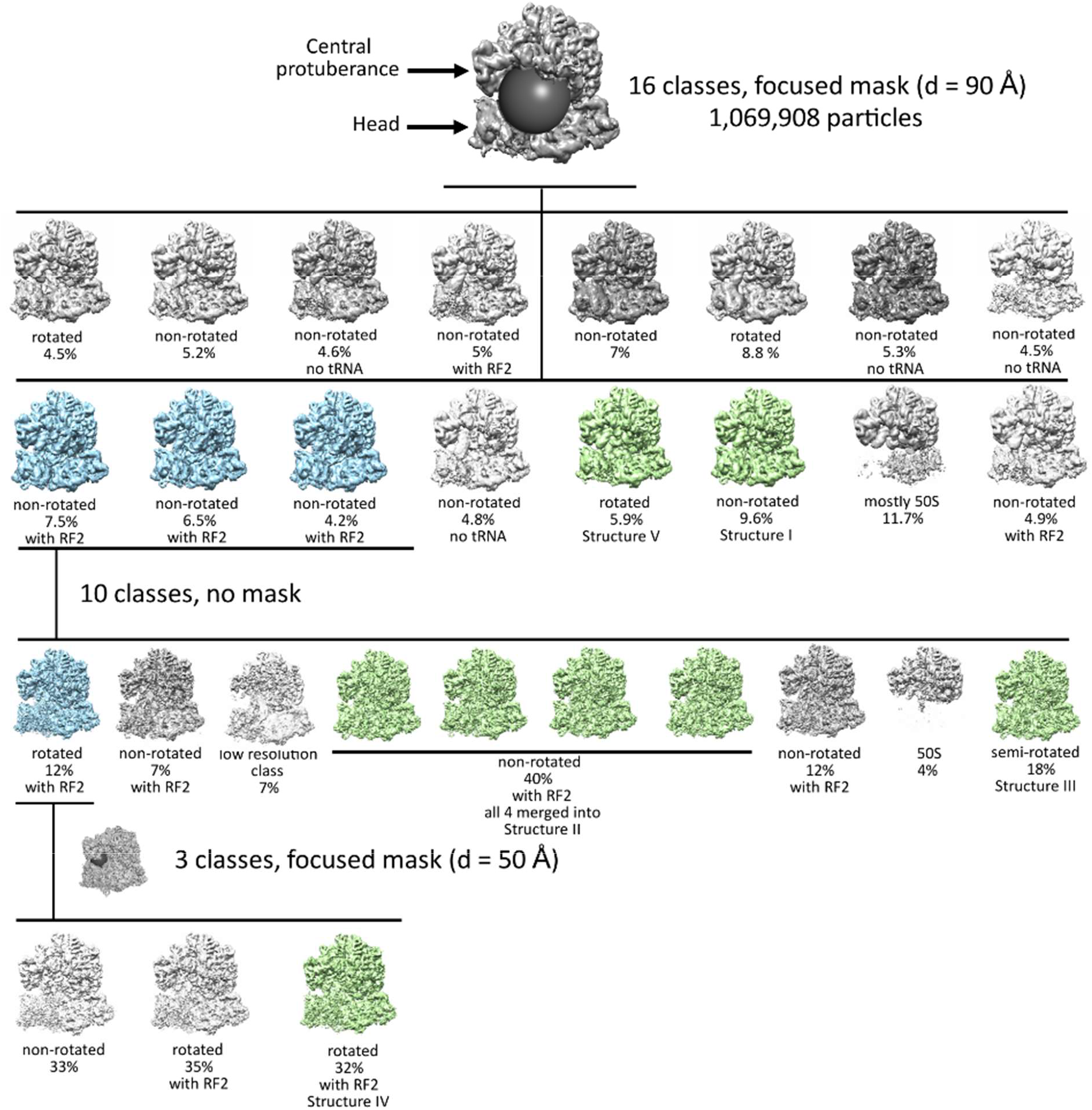
Maximum-likelihood classification strategies to obtain the final maps and state occupancies (shown as percentages of particles for each classification). Cyan classes were used for further classifications, green classes were used for obtaining the final maps. The views show the 50S (top) and 30S (bottom) subunits and the positions of spherical masks.

**Figure 1—figure supplement 2.**
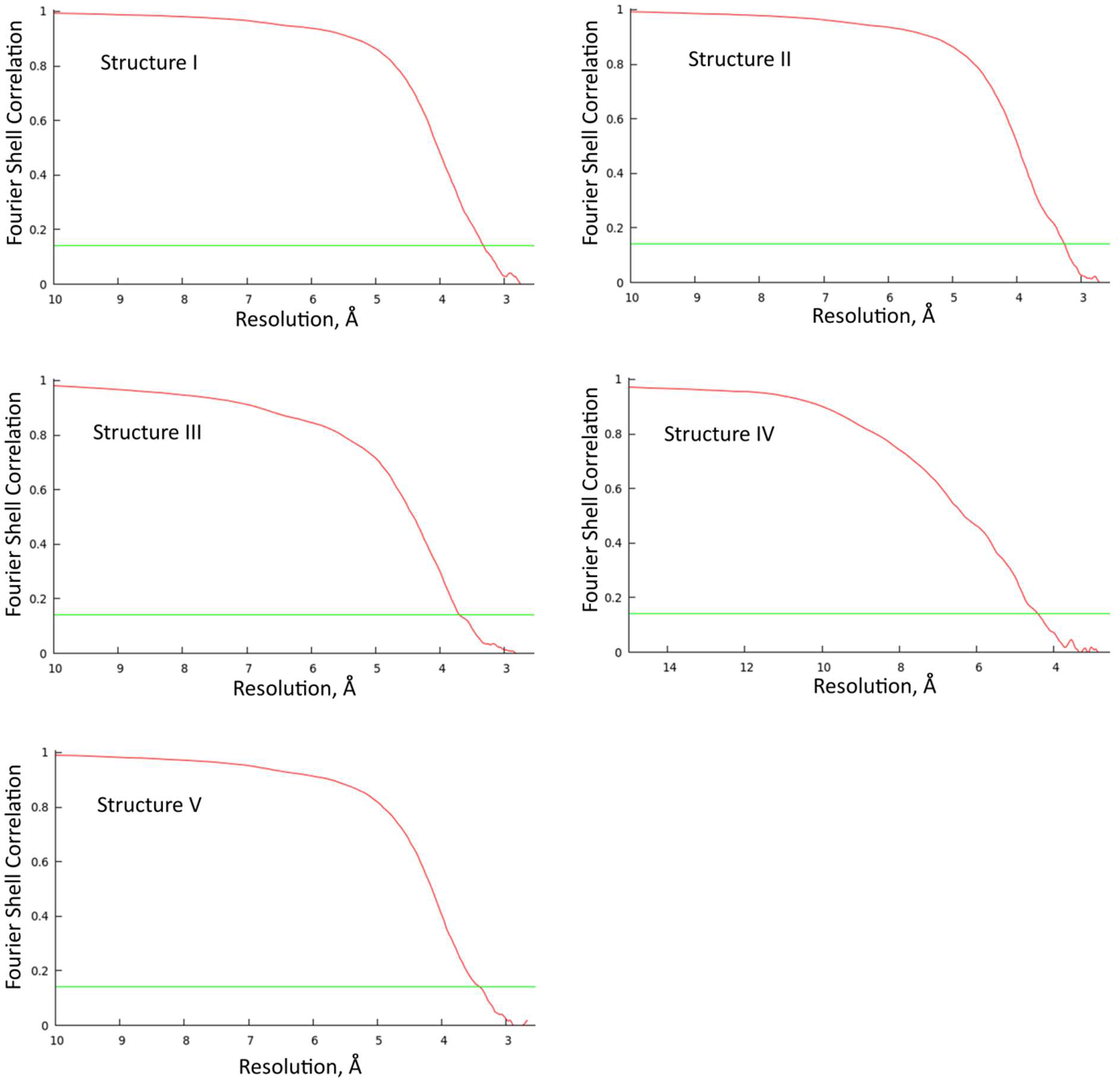
Fourier shell correlation (FSC) between even- and odd-particle half maps show that resolution ranges from 3.3 to 4.2 Å for Structures I-V (at FSC=0.143, green line).

**Figure 1—figure supplement 3.**
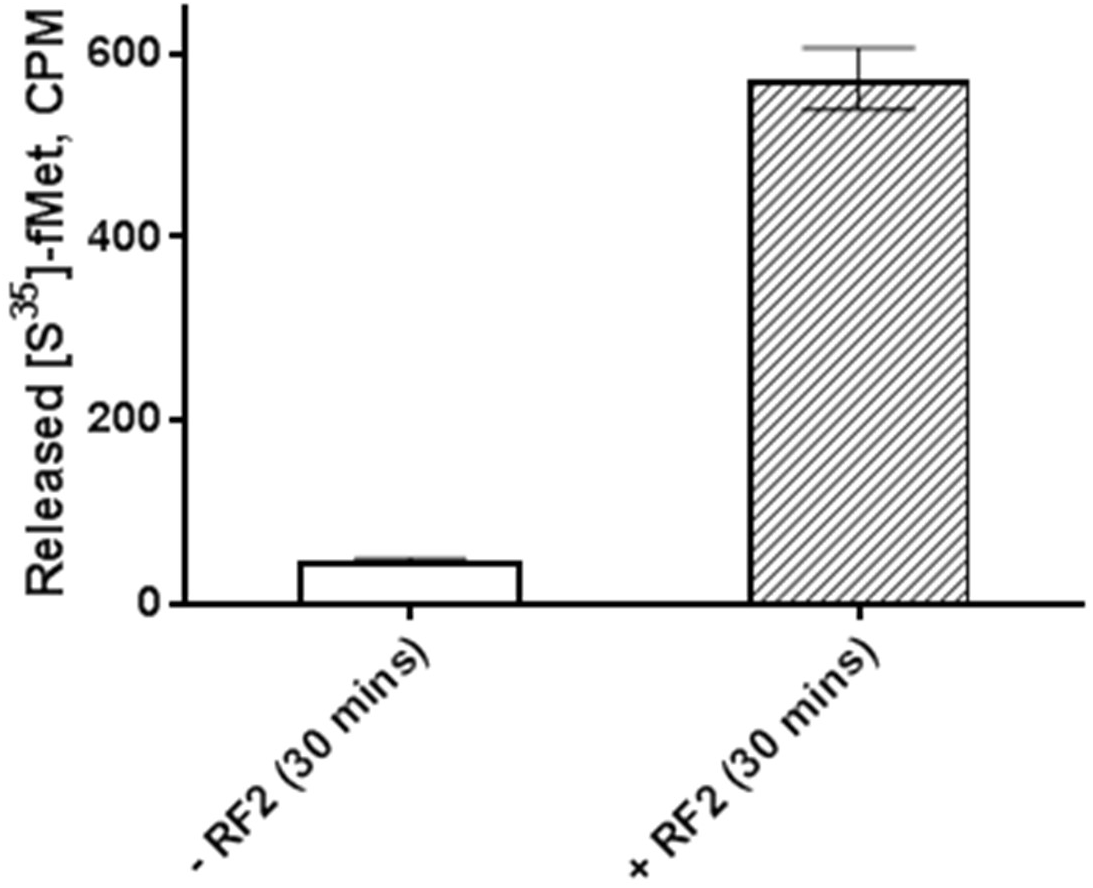
RF2 mediates formyl-methionine release from 70S ribosomes. Activity of RF2 is shown as counts from [S^35^]-fMet released from 70S•[S^35^]-fMet-tRNA^fMet^•mRNA(UGA), prepared using buffer conditions and concentrations identical to those used for cryo-EM grids preparation (see Methods). Column height represents an average of two measurements, error bars represent a standard error.

**Figure 2—figure supplement 1.**
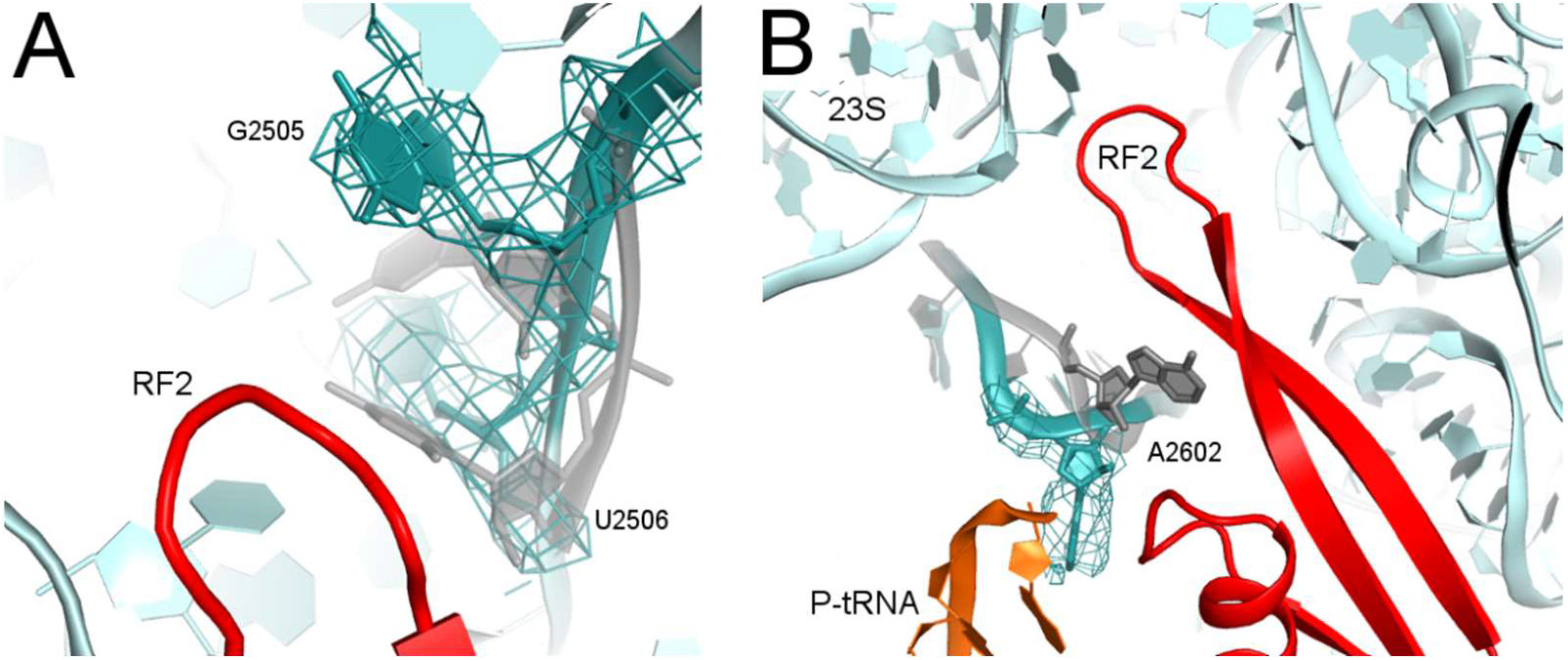
Rearrangements of PTC nucleotides (cyan) upon formation of the β-hairpin structure by the catalytic region of RF2 (red; Structure II is shown). Nucleotides in the crystal structure with an α-helical conformation of the GGQ motif of RF2 (not shown; see Figure 2A) are shown in gray (Korostelev et al., 2008). (A) Cryo-EM density (gray mesh, shown at σ=3.0) for G2505 and U2506 (shown as sticks). (B) Cryo-EM density (gray mesh, shown at σ=2.5) for A2602 (shown as sticks).

**Figure 3—figure supplement 1.**
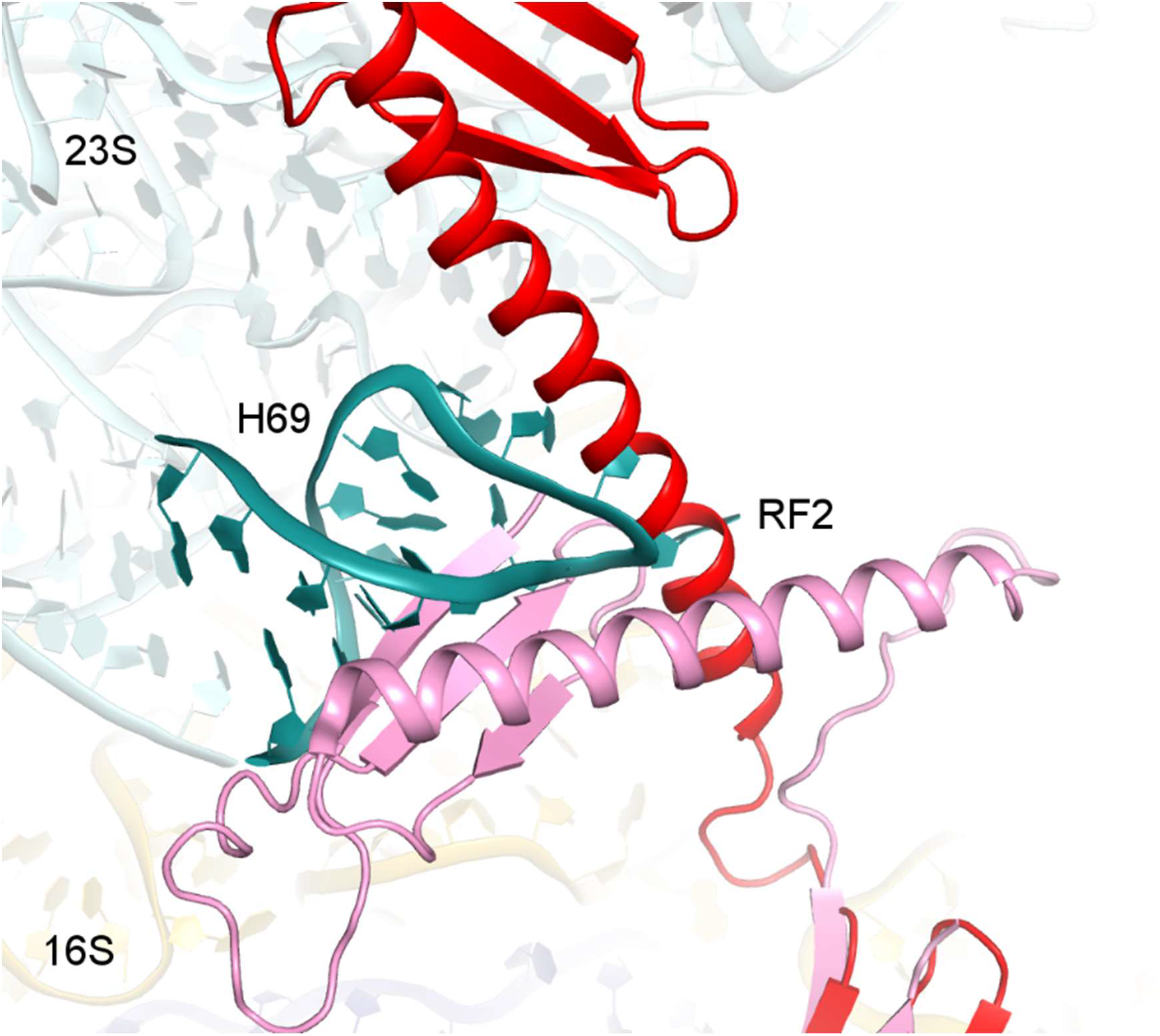
Structure alignments show that the disengaged H69 in Structure V (teal) is incompatible with the extended (red, Structure II) and compact (pink, PDB 1GQE (Vestergaard et al., 2001)) conformations of RF2 due to steric hindrance. Structures II and V were superimposed via 16S rRNA. Codon-recognition domain 2 of compact RF2 was superimposed with domain 2 of RF2 in Structure II.

**Movie 1. An animation showing transitions between the structures of the 70S ribosome upon interaction with RF2.**

## Acknowledgements

We thank KangKang Song for help with screening cryo-EM grids and for assistance with data collection at the UMass Medical School cryo-EM facility; Darryl Conte Jr. and members of the Korostelev lab for helpful comments on the manuscript. This study was supported by NIH Grants R01 GM107465 and R35 GM127094.

## Conflict of interest

The authors declare that they have no conflicts of interest with the contents of this article.

## Author contributions

E.S. assembled the ribosome complexes, performed biochemistry, collected and processed cryo-EM data, modeled and refined structural models; G.D. purified ribosome subunits and RF2, and assisted with structure refinements; C.X. implemented multiple-shot (per hole) data acquisition and optimized data collection; A.B.L assisted with data processing; A.A.K assisted with modeling and structure refinements, and wrote the manuscript with input from co-authors. All authors finalized the manuscript.

## Methods

### Preparation of the 70S termination complex with RF2

N-terminally His6-tagged release factor 2 (RF2) from *E. coli* K12 strain was overexpressed in *E. Coli* BLR (DE3) (Novagen) and purified as described for RF1 (Svidritskiy and Korostelev, 2018a). *E. coli* tRNA^fMet^ (Chemical Block) was aminoacylated as described (Lancaster and Noller, 2005). 70S ribosomes were prepared from *E. coli* (MRE600) as described (Svidritskiy and Korostelev, 2018a). Ribosomes were stored in the ribosome-storage buffer (20 mM Tris-HCl, pH 7.0; 100 mM NH_4_Cl; 12.5 mM MgCl2; 0.5 mM EDTA; 6 mM β-mercaptoethanol) at – 80°C. A model mRNA fragment, containing the Shine-Dalgarno sequence and a spacer to position the AUG codon in the P site and the UGA stop codon in the A site (GGC AAG GAG GUA AAA <u>AUG UGA</u> AAAAAA), was synthesized by IDT.

The 70S•mRNA•fMet-tRNA^fMet^ complex with RF2 was assembled *in vitro* by mixing 400 nM 70S ribosome (all concentrations are final) with 12 μM mRNA in buffer containing 20 mM Tris-acetate (pH 7.0), 100 mM NH_4_OAc, 15 mM Mg(OAc)_2_. After incubation for two minutes at 37°C, 600 nM fMet-tRNA^fMet^ was added and incubated for five minutes at 37°C. The Mg(OAc)_2_ concentration was adjusted to 10 mM by adding an equal volume of buffer (20 mM Tris-acetate (pH 7.0), 100 mM NH_4_OAc, 5 mM Mg(OAc)_2_) at room temperature. The release reaction was started by adding an equal volume of 4 μM RF2 (in 20 mM Tris, 100 mM NH_4_OAc, 10 mM Mg(OAc)_2_), yielding the following final concentrations: 0.2 μM 70S, 6 μM mRNA, 0.3 μM fMet-tRNA^fMet^, and 2 μM RF2. The solution was incubated for 30 minutes at room temperature prior to grid preparation and plunging.

The activity of *E. coli* RF2 was tested in an *in vitro* assay that measures the release of formyl-methionine from the 70S ribosome bound with [^35^S]-fMet-tRNA^fMet^ in the P site, as described (Svidritskiy and Korostelev, 2018a). Pre-termination complex was prepared using 0.2 μM *E. coli* 70S ribosomes (all concentrations are for the final reaction mixture upon initiation of the release reaction), 6 μM mRNA and 0.3 μM [^35^S]-fMet-tRNA^fMet^ ([^35^S]-methionine from Perkin Elmer). An aliquot (4.5 μl) of the pre-termination complex was quenched in 30 μl of 0.1 M HCl to represent the zero-time point. Pre-termination complex was split into two tubes. 5 μl of 20 μM RF2 was added to 45 μl of the pre-termination complex in one tube, yielding 2 μM RF2 in the reaction. Buffer without RF2 was added to the second tube to test spontaneous (RF2-independent) hydrolysis of [^35^S]-fMet-tRNA^fMet^. After 30 minutes, 5-μl aliquots were quenched in 30 μl of 0.1 M HCl. Samples were extracted with ethylacetate, and the amount of released [^35^S]-labeled N-formyl-methionine was measured using a scintillation counter (Beckman Coulter, Inc.) in 3.5 ml of Econo-Safe scintillation cocktail (RPI). All measurements were performed twice (**Figure 1— figure supplement 3**).

### Cryo-EM and image processing

Holey-carbon grids (QUANTIFOIL R 2/1, Cu 200, Quantifoil Micro Tools) were coated with carbon and glow discharged with 20 mA with negative polarity for 60 seconds in a PELCO easiGlow glow discharge unit. 2.8 μl of the 70S•mRNA•fMet-tRNA^fMet^•RF2 complex was applied to the grids. Grids were blotted at blotting force 10 for 4 s at room temperature, 100% humidity, and plunged into liquid ethane using a Vitrobot MK4 (FEI). Grids were stored in liquid nitrogen.

A dataset of 1,069,908 particles was collected on a Titan Krios (FEI) microscope (operating at 300 kV) equipped with a K2 Summit camera system (Gatan), with −0.5 to −1.8 μm defocus. Multi-shot data collection was performed by recording 6 exposures per hole, using SerialEM (Mastronarde, 2005) with beam-image shift. Coma-free alignment was performed using a built-in function. “Coma vs. Image Shift” from the Calibration menu was used for dynamic beam-tilt compensation, based on image shifts for each exposure. Multi-hole/multi-shot configuration was selected from “Multiple Record Setup Dialog” to dynamically adjust the beam tilt. Backlash-corrected compensation was applied to each stage movement at the target stage position to reduce mechanical stage drift. 7,287 movies were collected. Each exposure (32 frames per movie) was acquired with continuous frame streaming, yielding a total dose of 29.4 e^-^/Å^2^. The nominal magnification was 105,000 and the calibrated super-resolution pixel size at the specimen level was 0.667 Å. 6,972 movies were selected after discarding those with defects due to ice or image recording. The frames for each movie were processed using IMOD (Kremer et al., 1996) and binned to pixel size 1.334 Å (termed unbinned or 1× binned). Movies were motion-corrected, and frame averages were calculated using all 32 frames within each movie, after multiplying by the corresponding gain reference. cisTEM (Grant et al., 2018) was used to determine defocus values for each resulting frame average and for particle picking. The stack and particle parameter files were assembled in cisTEM with the binnings of 1×, 2×, 4×, and 6× (box size of 624 for a non-binned stack).

Data were processed essentially as described in our recent work (Loveland et al., 2017) and shown in **Figure 1—figure supplement 1**. FrealignX was used for particle alignment, refinement, and final reconstruction steps, and Frealign v9.11 was used for classification steps (Lyumkis et al., 2013). The 6×-binned image stack (1,069,908 particles) was initially aligned to a ribosome reference without release factor and tRNA (PDB 5J4D) (Svidritskiy et al., 2016) using 3 cycles of mode 3 (global search) alignment, including data from 20-Å to 300-Å resolution. Subsequently, the 6×-binned stack was refined using mode 1 (3 cycles) to align particles in the 15- to 300-Å resolution range. Using the 4×-binned image stack, the particles were successively refined using mode 1 by gradually increasing the high-resolution limit to 12- and 10-Å (3 cycles for each resolution limit). 3D density reconstruction was obtained using 60% of particles with highest scores. The map contained density for the P-tRNA, mRNA, and RF2.

Data classification was performed using a spherical focused mask with a 45-Å radius, centered roughly between the A-site finger (nucleotides 870–910 of 23S rRNA), protein L11, and peptidyl transferase center. The mask covered most of the ribosomal A site and part of the P site. The particle stack was classified into 16 classes, using the high-resolution limit of 10 Å. The classification resulted in five RF2-containing classes, six classes with tRNA alone (two classes represented Structures I and V), three classes of the 70S ribosome without tRNA, and two classes of the 50S subunit alone. All classes were subsequently reconstructed using the unbinned stack. Three strongest classes with RF2 were merged and extracted from the unbinned stack, and subclassified into 10 classes without masking, using a high-resolution limit of 8 Å. This classification yielded Structure II, four maps that were merged to obtain Structure III, a map with a rotated 70S•RF2 conformation (used for further classification to obtain Structure IV), a lower-resolution map similar to Structure II, and three junk (poorly resolved) classes. The class corresponding to the rotated 70S•RF2 conformation was extracted from the unbinned substack and further classified into 3 classes using a 25-Å radius spherical-focused mask roughly centered at the elbow of tRNA (high-resolution limit at 12 Å), yielding Structure IV. All final classes were reconstructed in FrealignX (1×-binning) using 95% of particles with highest scores.

The resulting reconstructions varied from 3.3 Å (Structure II) to 4.2 (Structure IV) Å average resolution (Fourier Shell Correlation (FSC) = 0.143). Cryo-EM reconstructions were B-factor sharpened in bfactor.exe (part of Frealign distribution) using different B-factor values (suggested by bfactor.exe) and then used for model building and structure refinements. B-factor of −50 Å^2^ was also used for initial refinements and to visualize lower-resolution details. FSC curves were calculated by FrealignX for even and odd particle half-sets (**Figure 1 – figure supplement 2**).

### Model building and refinement

Cryo-EM structures of *E. coli* 70S•ArfA•RF2 complex (Demo et al., 2017) excluding ArfA, and the 70S•fMet-tRNA^fmet^ complex with EF-Tu ternary complex (Loveland et al., 2017) and improved structure of S2 (Loveland and Korostelev, 2017), excluding the ternary complex, were used to create initial models for structure refinement. Chimera (Pettersen et al., 2004) was used for fitting the 50S subunit, 30S head, 30S body, mRNA, tRNA, and RF2 domains. Domain 1 of RF2 was poorly resolved in the maps and was omitted from structural models. Local model fitting was performed in Pymol (DeLano, 2002).

The structural model was refined by real-space simulated-annealing refinement using atomic electron scattering factors in RSRef, as described (Svidritskiy et al., 2014) (Chapman, 1995; Korostelev et al., 2002). Secondary-structure restraints, comprising hydrogen-bonding restraints for ribosomal proteins and base-pairing restraints for RNA molecules, were employed as described (Korostelev et al., 2008). Refinement parameters, such as the relative weighting of stereochemical restraints and experimental energy term, were optimized to produce the stereochemically optimal models that closely agree with the corresponding maps. In the final stage, the structures were refined using Phenix (phenix.real_space_refine) (Adams et al., 2010), followed by a round of refinement in RSRef, applying harmonic restraints to preserve protein backbone geometry. B-factors were refined in Phenix. The refined structural models closely agree with the corresponding maps and have good stereochemical parameters, characterized by low deviation from ideal bond lengths and angles, low numbers of protein-backbone outliers and other robust structure-quality statistics, as shown in Table 2. Structure quality was validated using MolProbity (Chen et al., 2010).

Structure superpositions were performed in PyMOL. For structural comparisons, the distances and angles were calculated in Pymol and Chimera, respectively. To calculate an angle of the 30S subunit rotation between two 70S structures, the 23S rRNAs were aligned using PyMOL, and the angle between 16S rRNAs body domains (all nucleotide residues excluding 925-1390) was measured in Chimera. To calculate an angle of the 30S-head rotation (swivel) between two 70S structures, the 16S rRNAs of the 30S body were aligned using PyMOL, and the angle between the 16S rRNA residues 925-1390 was measured in Chimera. Ribosome conformations (30S body/subunit rotation and 30S head rotation) for Structures I through V were compared with the crystal structure of the 70S•RF2 complex formed with the UGA stop codon (Weixlbaumer et al., 2008).

The cryo-EM maps were deposited in EMDB (Structure I – EMD-20048, Structure II – EMD-20052, Structure III – EMD-20056, Structure IV – EMD-20057, Structure V – EMD-20058). The PDB models were deposited in RCSB (PDB codes 6OFX, 6OG7, 6OGF, 6OGG, 6OGI respectively). Figures were prepared in PyMOL.

## References

Abeyrathne, P.D., Koh, C.S., Grant, T., Grigorieff, N., and Korostelev, A.A. (2016). Ensemble cryo-EM uncovers inchworm-like translocation of a viral IRES through the ribosome. Elife 5.

Adams, P.D., Afonine, P.V., Bunkoczi, G., Chen, V.B., Davis, I.W., Echols, N., Headd, J.J., Hung, L.W., Kapral, G.J., Grosse-Kunstleve, R.W., McCoy, A.J., Moriarty, N.W., Oeffner, R., Read, R.J., Richardson, D.C., Richardson, J.S., Terwilliger, T.C., and Zwart, P.H. (2010). PHENIX: a comprehensive Python-based system for macromolecular structure solution. Acta Crystallogr D Biol Crystallogr 66, 213–221.

Adio, S., Sharma, H., Senyushkina, T., Karki, P., Maracci, C., Wohlgemuth, I., Holtkamp, W., Peske, F., and Rodnina, M.V. (2018). Dynamics of ribosomes and release factors during translation termination in E. coli. Elife 7.

Agirrezabala, X., Lei, J., Brunelle, J.L., Ortiz-Meoz, R.F., Green, R., and Frank, J. (2008). Visualization of the hybrid state of tRNA binding promoted by spontaneous ratcheting of the ribosome. Mol Cell 32, 190–197.

Agirrezabala, X., Liao, H.Y., Schreiner, E., Fu, J., Ortiz-Meoz, R.F., Schulten, K., Green, R., and Frank, J. (2012). Structural characterization of mRNA-tRNA translocation intermediates. Proc Natl Acad Sci U S A 109, 6094–6099.

Agrawal, R.K., Sharma, M.R., Kiel, M.C., Hirokawa, G., Booth, T.M., Spahn, C.M., Grassucci, R.A., Kaji, A., and Frank, J. (2004). Visualization of ribosome-recycling factor on the Escherichia coli 70S ribosome: functional implications. Proc Natl Acad Sci U S A 101, 8900–8905.

Agrawal, R.K., Spahn, C.M., Penczek, P., Grassucci, R.A., Nierhaus, K.H., and Frank, J. (2000). Visualization of tRNA movements on the Escherichia coli 70S ribosome during the elongation cycle. J Cell Biol 150, 447–460.

Amort, M., Wotzel, B., Bakowska-Zywicka, K., Erlacher, M.D., Micura, R., and Polacek, N. (2007). An intact ribose moiety at A2602 of 23S rRNA is key to trigger peptidyl-tRNA hydrolysis during translation termination. Nucleic Acids Res 35, 5130–5140.

Borovinskaya, M.A., Pai, R.D., Zhang, W., Schuwirth, B.S., Holton, J.M., Hirokawa, G., Kaji, H., Kaji, A., and Cate, J.H. (2007). Structural basis for aminoglycoside inhibition of bacterial ribosome recycling. Nat Struct Mol Biol 14, 727–732.

Brenner, S., Barnett, L., Katz, E.R., and Crick, F.H. (1967). UGA: a third nonsense triplet in the genetic code. Nature 213, 449–450.

Brenner, S., Stretton, A.O., and Kaplan, S. (1965). Genetic code: the ‘nonsense’ triplets for chain termination and their suppression. Nature 206, 994–998.

Brilot, A.F., Korostelev, A.A., Ermolenko, D.N., and Grigorieff, N. (2013). Structure of the ribosome with elongation factor G trapped in the pretranslocation state. Proc Natl Acad Sci U S A.

Capecchi, M.R. (1967a). Polypeptide chain termination in vitro: isolation of a release factor. Proc Natl Acad Sci U S A 58, 1144–1151.

Capecchi, M.R. (1967b). A rapid assay for polypeptide chain termination. Biochem Biophys Res Commun 28, 773–778.

Casy, W., Prater, A.R., and Cornish, P.V. (2018). Operative Binding of Class I Release Factors and YaeJ Stabilizes the Ribosome in the Nonrotated State. Biochemistry 57, 1954–1966.

Chapman, M.S. (1995). Restrained real-space macromolecular atomic refinement using a new resolution-dependent electron density function. Acta Crystallogr. A 51, 69–80.

Chen, V.B., Arendall, W.B., 3rd, Headd, J.J., Keedy, D.A., Immormino, R.M., Kapral, G.J., Murray, L.W., Richardson, J.S., and Richardson, D.C. (2010). MolProbity: all-atom structure validation for macromolecular crystallography. Acta Crystallogr D Biol Crystallogr 66, 12–21.

Chen, Y., Feng, S., Kumar, V., Ero, R., and Gao, Y.G. (2013). Structure of EF-G-ribosome complex in a pretranslocation state. Nat Struct Mol Biol 20, 1077–1084.

Cornish, P.V., Ermolenko, D.N., Noller, H.F., and Ha, T. (2008). Spontaneous intersubunit rotation in single ribosomes. Mol Cell 30, 578–588.

DeLano, W.L. (2002). The PyMOL Molecular Graphics System. (Palo Alto, CA, USA.: DeLano Scientific).

Demo, G., Svidritskiy, E., Madireddy, R., Diaz-Avalos, R., Grant, T., Grigorieff, N., Sousa, D., and Korostelev, A.A. (2017). Mechanism of ribosome rescue by ArfA and RF2. Elife 6.

Dunkle, J.A., and Cate, J.H. (2010). Ribosome structure and dynamics during translocation and termination. Annu Rev Biophys 39, 227–244.

Dunkle, J.A., Wang, L., Feldman, M.B., Pulk, A., Chen, V.B., Kapral, G.J., Noeske, J., Richardson, J.S., Blanchard, S.C., and Cate, J.H. (2011). Structures of the bacterial ribosome in classical and hybrid states of tRNA binding. Science 332, 981–984.

Ermolenko, D.N., Majumdar, Z.K., Hickerson, R.P., Spiegel, P.C., Clegg, R.M., and Noller, H.F. (2007). Observation of intersubunit movement of the ribosome in solution using FRET. J Mol Biol 370, 530–540.

Fischer, N., Konevega, A.L., Wintermeyer, W., Rodnina, M.V., and Stark, H. (2010). Ribosome dynamics and tRNA movement by time-resolved electron cryomicroscopy. Nature 466, 329–333.

Florin, T., Maracci, C., Graf, M., Karki, P., Klepacki, D., Berninghausen, O., Beckmann, R., Vazquez-Laslop, N., Wilson, D.N., Rodnina, M.V., and Mankin, A.S. (2017). An antimicrobial peptide that inhibits translation by trapping release factors on the ribosome. Nat Struct Mol Biol 24, 752–757.

Frank, J., and Agrawal, R.K. (2000). A ratchet-like inter-subunit reorganization of the ribosome during translocation. Nature 406, 318–322.

Freistroffer, D.V., Pavlov, M.Y., MacDougall, J., Buckingham, R.H., and Ehrenberg, M. (1997). Release factor RF3 in E.coli accelerates the dissociation of release factors RF1 and RF2 from the ribosome in a GTP-dependent manner. EMBO J 16, 4126–4133.

Fu, Z., Indrisiunaite, G., Kaledhonkar, S., Shah, B., Sun, M., Chen, B., Grassucci, R.A., Ehrenberg, M., and Frank, J. (2018). The structural basis for release factor activation during translation termination revealed by time-resolved cryogenic electron microscopy. bioRxiv November 4.

Fu, Z., Kaledhonkar, S., Borg, A., Sun, M., Chen, B., Grassucci, R.A., Ehrenberg, M., and Frank, J. (2016). Key Intermediates in Ribosome Recycling Visualized by Time-Resolved Cryoelectron Microscopy. Structure 24, 2092–2101.

Gao, H., Zhou, Z., Rawat, U., Huang, C., Bouakaz, L., Wang, C., Cheng, Z., Liu, Y., Zavialov, A., Gursky, R., Sanyal, S., Ehrenberg, M., Frank, J., and Song, H. (2007). RF3 induces ribosomal conformational changes responsible for dissociation of class I release factors. Cell 129, 929–941.

Gao, Y.G., Selmer, M., Dunham, C.M., Weixlbaumer, A., Kelley, A.C., and Ramakrishnan, V. (2009). The structure of the ribosome with elongation factor G trapped in the posttranslocational state. Science 326, 694–699.

Graf, M., Huter, P., Maracci, C., Peterek, M., Rodnina, M.V., and Wilson, D.N. (2018). Visualization of translation termination intermediates trapped by the Apidaecin 137 peptide during RF3-mediated recycling of RF1. Nat Commun 9, 3053.

Grant, T., Rohou, A., and Grigorieff, N. (2018). cisTEM, user-friendly software for single-particle image processing. Elife 7.

Grentzmann, G., Brechemier-Baey, D., Heurgue, V., Mora, L., and Buckingham, R.H. (1994). Localization and characterization of the gene encoding release factor RF3 in Escherichia coli. Proc Natl Acad Sci U S A 91, 5848–5852.

He, S.L., and Green, R. (2010). Visualization of codon-dependent conformational rearrangements during translation termination. Nat Struct Mol Biol 17, 465–470.

Hetrick, B., Lee, K., and Joseph, S. (2009). Kinetics of stop codon recognition by release factor 1. Biochemistry 48, 11178–11184.

James, N.R., Brown, A., Gordiyenko, Y., and Ramakrishnan, V. (2016). Translational termination without a stop codon. Science 354, 1437–1440.

JCSG (2005). Crystal structure of Peptide chain release factor 1 (RF-1) (SMU.1085) from Streptococcus mutans at 2.34 A resolution. J.C.f.S. Genomics, ed. (Protein Data Bank).

Jin, H., Kelley, A.C., Loakes, D., and Ramakrishnan, V. (2010). Structure of the 70S ribosome bound to release factor 2 and a substrate analog provides insights into catalysis of peptide release. Proc Natl Acad Sci U S A 107, 8593–8598.

Jin, H., Kelley, A.C., and Ramakrishnan, V. (2011). Crystal structure of the hybrid state of ribosome in complex with the guanosine triphosphatase release factor 3. Proc Natl Acad Sci U S A 108, 15798–15803.

Julian, P., Konevega, A.L., Scheres, S.H., Lazaro, M., Gil, D., Wintermeyer, W., Rodnina, M.V., and Valle, M. (2008). Structure of ratcheted ribosomes with tRNAs in hybrid states. Proc Natl Acad Sci U S A 105, 16924–16927.

Klaholz, B.P., Pape, T., Zavialov, A.V., Myasnikov, A.G., Orlova, E.V., Vestergaard, B., Ehrenberg, M., and van Heel, M. (2003). Structure of the Escherichia coli ribosomal termination complex with release factor 2. Nature 421, 90–94.

Korostelev, A., Asahara, H., Lancaster, L., Laurberg, M., Hirschi, A., Zhu, J., Trakhanov, S., Scott, W.G., and Noller, H.F. (2008). Crystal structure of a translation termination complex formed with release factor RF2. Proc Natl Acad Sci U S A 105, 19684–19689.

Korostelev, A., Bertram, R., and Chapman, M.S. (2002). Simulated-annealing real-space refinement as a tool in model building. Acta Crystallogr D Biol Crystallogr 58, 761–767.

Korostelev, A., Trakhanov, S., Laurberg, M., and Noller, H.F. (2006). Crystal structure of a 70S ribosome-tRNA complex reveals functional interactions and rearrangements. Cell 126, 1065–1077.

Korostelev, A., Zhu, J., Asahara, H., and Noller, H.F. (2010). Recognition of the amber UAG stop codon by release factor RF1. EMBO J 29, 2577–2585.

Korostelev, A.A. (2011). Structural aspects of translation termination on the ribosome. RNA 17, 1409–1421.

Koutmou, K.S., McDonald, M.E., Brunelle, J.L., and Green, R. (2014). RF3:GTP promotes rapid dissociation of the class 1 termination factor. RNA 20, 609–620.

Kremer, J.R., Mastronarde, D.N., and McIntosh, J.R. (1996). Computer visualization of three-dimensional image data using IMOD. J Struct Biol 116, 71–76.

Lancaster, L., and Noller, H.F. (2005). Involvement of 16S rRNA nucleotides G1338 and A1339 in discrimination of initiator tRNA. Mol Cell 20, 623–632.

Laurberg, M., Asahara, H., Korostelev, A., Zhu, J., Trakhanov, S., and Noller, H.F. (2008). Structural basis for translation termination on the 70S ribosome. Nature 454, 852–857.

Leipe, D.D., Wolf, Y.I., Koonin, E.V., and Aravind, L. (2002). Classification and evolution of P-loop GTPases and related ATPases. J Mol Biol 317, 41–72.

Ling, C., and Ermolenko, D.N. (2016). Structural insights into ribosome translocation. WIREs RNA.

Loveland, A.B., Demo, G., Grigorieff, N., and Korostelev, A.A. (2017). Ensemble cryo-EM elucidates the mechanism of translation fidelity. Nature 546, 113–117.

Loveland, A.B., and Korostelev, A.A. (2017). Structural dynamics of protein S1 on the 70S ribosome visualized by ensemble cryo-EM. Methods.

Lyumkis, D., Brilot, A.F., Theobald, D.L., and Grigorieff, N. (2013). Likelihood-based classification of cryo-EM images using FREALIGN. J Struct Biol 183, 377–388.

Mastronarde, D.N. (2005). Automated electron microscope tomography using robust prediction of specimen movements. J Struct Biol 152, 36–51.

Mikuni, O., Ito, K., Moffat, J., Matsumura, K., McCaughan, K., Nobukuni, T., Tate, W., and Nakamura, Y. (1994). Identification of the prfC gene, which encodes peptide-chain-release factor 3 of Escherichia coli. Proc Natl Acad Sci U S A 91, 5798–5802.

Moazed, D., and Noller, H.F. (1989). Intermediate states in the movement of transfer RNA in the ribosome. Nature 342, 142–148.

Nakamura, Y., Ito, K., Matsumura, K., Kawazu, Y., and Ebihara, K. (1995). Regulation of translation termination: conserved structural motifs in bacterial and eukaryotic polypeptide release factors. Biochem Cell Biol 73, 1113–1122.

Nichols, R.J., Sen, S., Choo, Y.J., Beltrao, P., Zietek, M., Chaba, R., Lee, S., Kazmierczak, K.M., Lee, K.J., Wong, A., Shales, M., Lovett, S., Winkler, M.E., Krogan, N.J., Typas, A., and Gross, C.A. (2011). Phenotypic landscape of a bacterial cell. Cell 144, 143–156.

Noller, H.F., Lancaster, L., Zhou, J., and Mohan, S. (2017). The ribosome moves: RNA mechanics and translocation. Nat Struct Mol Biol 24, 1021–1027.

O’Connor, M. (2015). Interactions of release factor RF3 with the translation machinery. Mol Genet Genomics 290, 1335–1344.

Pai, R.D., Zhang, W., Schuwirth, B.S., Hirokawa, G., Kaji, H., Kaji, A., and Cate, J.H. (2008). Structural Insights into ribosome recycling factor interactions with the 70S ribosome. J Mol Biol 376, 1334–1347.

Petry, S., Brodersen, D.E., Murphy, F.V.t., Dunham, C.M., Selmer, M., Tarry, M.J., Kelley, A.C., and Ramakrishnan, V. (2005). Crystal structures of the ribosome in complex with release factors RF1 and RF2 bound to a cognate stop codon. Cell 123, 1255–1266.

Pettersen, E.F., Goddard, T.D., Huang, C.C., Couch, G.S., Greenblatt, D.M., Meng, E.C., and Ferrin, T.E. (2004). UCSF Chimera--a visualization system for exploratory research and analysis. J Comput Chem 25, 1605–1612.

Pierson, W.E., Hoffer, E.D., Keedy, H.E., Simms, C.L., Dunham, C.M., and Zaher, H.S. (2016). Uniformity of Peptide Release Is Maintained by Methylation of Release Factors. Cell Rep 17, 11–18.

Polacek, N., Gomez, M.J., Ito, K., Xiong, L., Nakamura, Y., and Mankin, A. (2003). The critical role of the universally conserved A2602 of 23S ribosomal RNA in the release of the nascent peptide during translation termination. Mol Cell 11, 103–112.

Prabhakar, A., Capece, M.C., Petrov, A., Choi, J., and Puglisi, J.D. (2017). Post-termination Ribosome Intermediate Acts as the Gateway to Ribosome Recycling. Cell Rep 20, 161–172.

Pulk, A., and Cate, J.H. (2013). Control of ribosomal subunit rotation by elongation factor G. Science 340, 1235970.

Ramakrishnan, V. (2011). Structural studies on decoding, termination and translocation in the bacterial ribosome In Ribosomes: Structure, Function, and Dynamics. M. Rodnina, W. Wintermeyer, and R. Green, eds. (Springer), pp. 19–30.

Ratje, A.H., Loerke, J., Mikolajka, A., Brunner, M., Hildebrand, P.W., Starosta, A.L., Donhofer, A., Connell, S.R., Fucini, P., Mielke, T., Whitford, P.C., Onuchic, J.N., Yu, Y., Sanbonmatsu, K.Y., Hartmann, R.K., Penczek, P.A., Wilson, D.N., and Spahn, C.M. (2010). Head swivel on the ribosome facilitates translocation by means of intra-subunit tRNA hybrid sites. Nature 468, 713–716.

Rawat, U., Gao, H., Zavialov, A., Gursky, R., Ehrenberg, M., and Frank, J. (2006). Interactions of the release factor RF1 with the ribosome as revealed by cryo-EM. J Mol Biol 357, 1144–1153.

Rawat, U.B., Zavialov, A.V., Sengupta, J., Valle, M., Grassucci, R.A., Linde, J., Vestergaard, B., Ehrenberg, M., and Frank, J. (2003). A cryo-electron microscopic study of ribosome-bound termination factor RF2. Nature 421, 87–90.

Rodnina, M.V. (2018). Translation in Prokaryotes. Cold Spring Harb Perspect Biol 10.

Santos, N., Zhu, J., Donohue, J.P., Korostelev, A.A., and Noller, H.F. (2013). Crystal structure of the 70S ribosome bound with the Q253P mutant form of release factor RF2. Structure 21, 1258–1263.

Scolnick, E., Tompkins, R., Caskey, T., and Nirenberg, M. (1968). Release factors differing in specificity for terminator codons. Proc Natl Acad Sci U S A 61, 768–774.

Selmer, M., Dunham, C.M., Murphy, F.V., Weixlbaumer, A., Petry, S., Kelley, A.C., Weir, J.R., and Ramakrishnan, V. (2006). Structure of the 70S ribosome complexed with mRNA and tRNA. Science 313, 1935–1942.

Shin, D.H., Brandsen, J., Jancarik, J., Yokota, H., Kim, R., and Kim, S.H. (2004). Structural analyses of peptide release factor 1 from Thermotoga maritima reveal domain flexibility required for its interaction with the ribosome. J Mol Biol 341, 227–239.

Sternberg, S.H., Fei, J., Prywes, N., McGrath, K.A., and Gonzalez, R.L., Jr. (2009). Translation factors direct intrinsic ribosome dynamics during translation termination and ribosome recycling. Nat Struct Mol Biol 16, 861–868.

Svidritskiy, E., Brilot, A.F., Koh, C.S., Grigorieff, N., and Korostelev, A.A. (2014). Structures of Yeast 80S Ribosome-tRNA Complexes in the Rotated and Nonrotated Conformations. Structure 22, 1210–1218.

Svidritskiy, E., Demo, G., and Korostelev, A.A. (2018). Mechanism of premature translation termination on a sense codon. J Biol Chem 293, 12472–12479.

Svidritskiy, E., and Korostelev, A.A. (2018a). Conformational control of translation termination on the 70S ribosome. Structure 26, 821–828.

Svidritskiy, E., and Korostelev, A.A. (2018b). Mechanism of Inhibition of Translation Termination by Blasticidin S. J Mol Biol 430, 591–593.

Svidritskiy, E., Madireddy, R., and Korostelev, A.A. (2016). Structural Basis for Translation Termination on a Pseudouridylated Stop Codon. J Mol Biol 428, 2228–2236.

Tompkins, R.K., Scolnick, E.M., and Caskey, C.T. (1970). Peptide chain termination. VII. The ribosomal and release factor requirements for peptide release. Proc Natl Acad Sci U S A 65, 702–708.

Tourigny, D.S., Fernandez, I.S., Kelley, A.C., and Ramakrishnan, V. (2013). Elongation factor G bound to the ribosome in an intermediate state of translocation. Science 340, 1235490.

Trappl, K., and Joseph, S. (2016). Ribosome Induces a Closed to Open Conformational Change in Release Factor 1. J Mol Biol 428, 1333–1344.

Vestergaard, B., Van, L.B., Andersen, G.R., Nyborg, J., Buckingham, R.H., and Kjeldgaard, M. (2001). Bacterial polypeptide release factor RF2 is structurally distinct from eukaryotic eRF1. Mol Cell 8, 1375–1382.

Weixlbaumer, A., Jin, H., Neubauer, C., Voorhees, R.M., Petry, S., Kelley, A.C., and Ramakrishnan, V. (2008). Insights into translational termination from the structure of RF2 bound to the ribosome. Science 322, 953–956.

Youngman, E.M., McDonald, M.E., and Green, R. (2008). Peptide release on the ribosome: mechanism and implications for translational control. Annu Rev Microbiol 62, 353–373.

Zeng, F., and Jin, H. (2018). Conformation of methylated GGQ in the Peptidyl Transferase Center during Translation Termination. Sci Rep 8, 2349.

Zhou, J., Lancaster, L., Donohue, J.P., and Noller, H.F. (2013). Crystal structures of EF-G-ribosome complexes trapped in intermediate states of translocation. Science 340, 1236086.

Zhou, J., Lancaster, L., Trakhanov, S., and Noller, H.F. (2012). Crystal structure of release factor RF3 trapped in the GTP state on a rotated conformation of the ribosome. RNA 18, 230–240.

Zoldak, G., Redecke, L., Svergun, D.I., Konarev, P.V., Voertler, C.S., Dobbek, H., Sedlak, E., and Sprinzl, M. (2007). Release factors 2 from Escherichia coli and Thermus thermophilus: structural, spectroscopic and microcalorimetric studies. Nucleic Acids Res 35, 1343–1353.

